# Natural H5N1 immunity in dairy cows is durable and cross-protective but non-sterilizing

**DOI:** 10.64898/2026.01.06.697911

**Authors:** Natalie N. Tarbuck, Hannah J. Cochran, Emma A. Martin, Min Liu, Kalyna S. Kulchytsky, William M. Leone, Richard J. Webby, Cody J. Warren, Andrew S. Bowman

**Author notes:** Corresponding Authors: Andrew Bowman,. Cody Warren.

## Abstract

Ongoing transmission of influenza A virus (H5N1) in U.S. dairy cattle threatens both animal and human health, underscoring the need to understand the durability of host immunity against reinfection with evolving genotypes. We challenged naïve and convalescent cows, infected one year prior with H5N1 genotype B3.13, with either homologous B3.13 or heterologous D1.1 genotype virus. Homologous rechallenge resulted in complete clinical protection with no infectious viral shedding. Conversely, heterologous rechallenge led to transient clinical disease and limited infectious viral shedding. Convalescent cows experienced significantly milder disease than naive cows, which developed severe illness with high viral shedding and required early euthanasia, regardless of the strain. These findings indicate that naturally acquired immunity offers strong protection against severe illness but may allow silent transmission of divergent strains. Therefore, natural herd immunity alone is unlikely to eliminate the virus; controlling H5N1 in cattle will likely require vaccination strategies that address viral evolution.

## INTRODUCTION

Influenza A viruses (IAVs) circulate widely among avian and mammalian hosts, exhibiting extraordinary genomic diversity and a well-documented capacity to cross species barriers. Since 2021, a global panzootic of highly pathogenic H5N1 clade 2.3.4.4b viruses has resulted in widespread infections across diverse mammalian species, including mink, skunks, raccoons, opossums, foxes, domestic cats, bears, and marine mammals^1–8^. In North America, reassortment between these 2.3.4.4b viruses and co-circulating low pathogenic avian influenza (LPAI) viruses has generated >100 distinct genotypes to date^9^, underscoring the extraordinary evolutionary plasticity of this lineage.

The recent spillover of IAV(H5N1) from wild birds into U.S. dairy cattle represents a major shift in the virus’s ecology, as cattle were not previously considered natural hosts for IAV. Yet, in less than a year, three independent introductions of IAV(H5N1) from wild birds have been detected in U.S. dairy cattle herds. This includes the detection of the B3.13 genotype in Texas in March 2024^10^, followed shortly thereafter by the emergence of the D1.1 genotype in Nevada (January 2025)^11^ and Arizona (February 2025)^12^. To date, more than 1,000 U.S. dairy cattle herds have been infected across 18 U.S. states^13^.While 41 human cases have been linked to direct exposure to cattle^14^, this is likely a substantial undercount given limited surveillance at the cattle-human interface.

IAV(H5N1) clade 2.3.4.4b infection in dairy cows exhibits an unprecedented tropism for the mammary gland with high viral titers detected in milk^4,15–18^. This is especially concerning given that milk is the economic foundation of the dairy industry and a major consumer commodity. Should IAV(H5N1) become endemic in dairy cows, it would potentially undermine a vital food animal industry. Further, continued circulation of IAV(H5N1) in dairy herds may lead to mammalian-adaptive mutations or reassortment with other IAV strains, potentially enhancing its ability to infect humans. At the same time, evolving genotypes in wild birds—with unknown potential to evade existing immunity in dairy cows—creates opportunities for additional introductions and persistence of the virus in dairy cows. Together, these concerns emphasize the ongoing and unprecedented threat that IAVs pose to the U.S. dairy industry and public health.

As the IAV(H5N1) epizootic persists into a second year, the durability and protective capacity of naturally acquired immunity in cattle remain undefined. This represents a critical gap in understanding the long-term consequences of continued IAV(H5N1) spread for both herd and human health risk. The B3.13 genotype viruses circulated widely in dairy cows following its initial introduction, and several months later D1.1 genotype viruses were also detected. It is unclear whether prior infection with one genotype provides cross-protection against a heterologous genotype, and to what extent naturally acquired immunity influences both clinical outcomes and viral replication. To assess these unknowns, dairy cows that had recovered from natural B3.13 infection one year earlier were experimentally rechallenged via the intramammary route with either the homologous B3.13 or heterologous D1.1 genotype viruses. We demonstrate that naturally acquired immunity in dairy cows provides robust protection against homologous and partial protection against heterologous IAV(H5N1) rechallenge one year after initial on farm exposure. Our findings provide important insights for future circulation of IAV(H5N1) in dairy cattle, as well as rationale for vaccination as a nationwide prevention strategy.

## RESULTS

### Detectable serum and milk antibody titers in cows naturally exposed to IAV(H5N1) B3.13 genotype more than one year later

After the detection of the IAV(H5N1) B3.13 genotype in Texas dairy cows in March 2024, the virus was introduced into an Ohio dairy herd through interstate cattle transport. Within this Ohio herd, approximately 20% of the cows showed clinical signs over three weeks, including reduced milk production, decreased rumination, and mastitis, resulting in significant economic losses. Serologic surveillance indicated nearly 90% of the herd seroconverted, revealing rapid and widespread transmission of the virus on farm^17^. Additional individual-level diagnostic testing (real time reverse transcriptase PCR, rRT-PCR) was performed on milk samples from selected clinically ill animals, confirming active IAV(H5N1) infections.

One year after the outbreak, we identified eight cows still in the Ohio herd that were clinically ill during the outbreak and diagnosed with IAV(H5N1) through rRT-PCR testing of their milk samples on April 5, 2024 (**Supplemental Table 1**). One year after the outbreak, we assessed the presence and magnitude of antibody responses in serum and milk from these animals prior to virus challenge. All previously infected cows had measurable serum antibody levels above the detection threshold in both hemagglutination-inhibition (HAI) assays performed using an attenuated IAV(H5N1) isolate and virus-neutralization (VNT) assays using the homologous IAV(H5N1) B3.13 genotype virus (**Figure 1A**). We next assayed for the presence of neutralizing antibodies in milk. Since the bovine udder comprises four mammary glands—also referred to as quarters—milk antibody responses were assessed separately for each quarter (left front (LF), right front (RF), left rear (LR), and right rear (RR)). In previously infected animals, neutralizing antibody levels were also detectable in milk, with titers varying by quarter within individual cows. The quarter-specific variability highlights a heterogeneous immune response across the udder and the compartmentalized nature of the four mammary glands. Of note, on-farm diagnostic testing during the initial outbreak was performed on composite milk samples representing each quarter, preventing determination of which mammary quarter(s) were originally infected. However, variation in antibody levels across the udder would be expected if these animals were not uniformly exposed to virus in each quarter.

**Figure 1.**
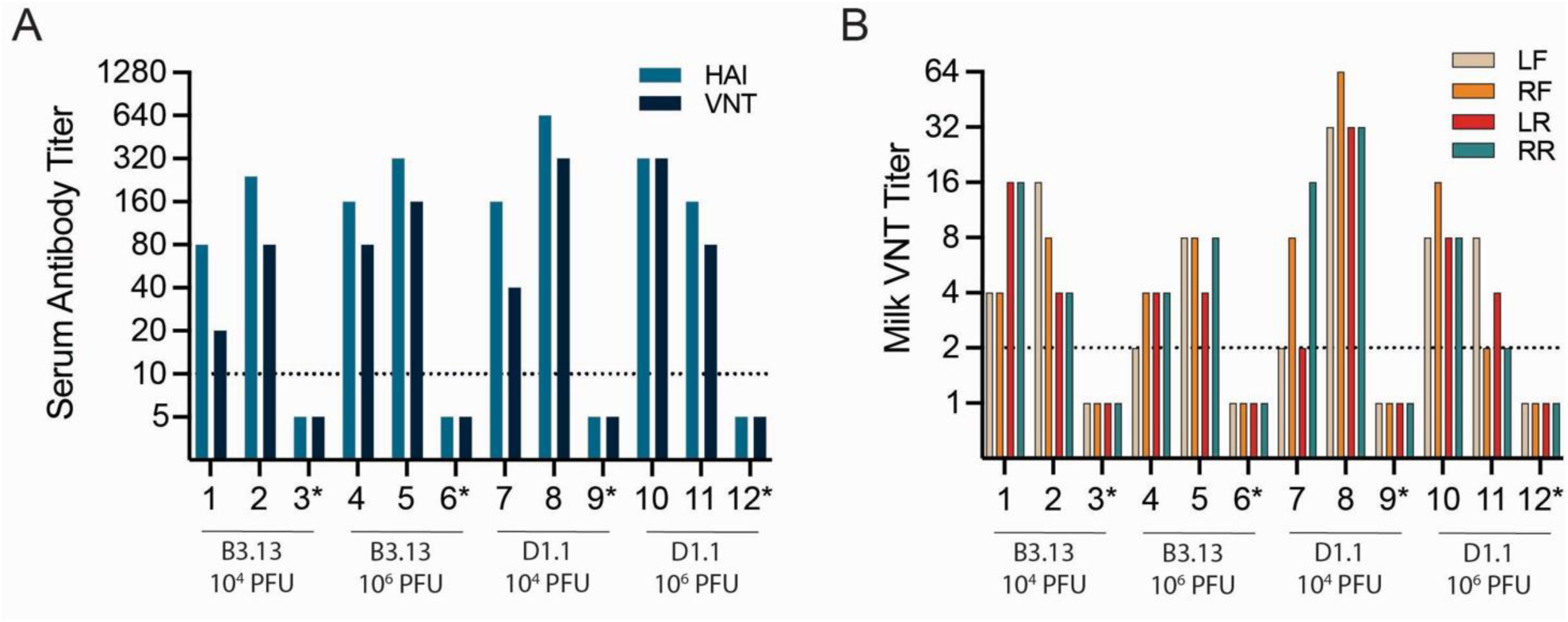
Detectable serum and milk antibody titers in cows naturally exposed to IAV(H5N1) B3.13 genotype more than one year later. Antibody titers were determined from samples collected prior to virus challenge. Cows were assigned to four infection groups to be challenged with either the B3.13 or D1.1 genotype virus at 10^4^ or 10^6^ PFU/cow. Each group contained two cows with natural immunity and one naïve control cow (controls indicated with an asterisk (*)). (**A**) Serum antibody titers were measured using hemagglutination inhibition (HAI) and virus neutralization (VNT) assays. (**B**) Milk virus neutralization titers are shown separately for each udder quarter: left front (LF), right front (RF), left rear (LR), and right rear (RR). The dotted line denotes the assay limit of detection. Naïve control cows exhibiting no response in either assay were set at half the limit of detection to indicate non-detectable antibody titers.

For comparison, four naïve control cows were sourced from a separate herd with no documented exposure to IAV(H5N1). Control cows lacked detectable neutralizing antibody responses in both serum and milk (denoted with “*” in **Figure 1A** and **1B**), confirming that they had not been previously exposed to IAV(H5N1). Collectively, these findings indicate natural immunity persists for more than one year after natural infection with IAV(H5N1) B3.13 genotype. Further, our results provide a rationale for examining the durability of immunity following re-exposure to IAV(H5N1).

### Naturally acquired IAV(H5N1) immunity protects against severe morbidity and mortality following subsequent challenge

Here, we evaluated whether previous infection with IAV(H5N1) B3.13 genotype protected cows from clinical disease upon rechallenge with a homologous or heterologous virus. We used two infectious doses (10^4^ PFU or 10^6^ PFU per cow) to evaluate if prior immunity modulates disease severity in a dose-dependent manner. The cows were assigned to four groups receiving either B3.13 or D1.1 virus at 10^4^ or 10^6^ PFU per cow, with each group containing two previously infected animals and one naive control (**Supplemental Figure 1)**. Viral stocks were resuspended in 10 mL of infection media and distributed evenly across all four quarters (2.5 mL per teat canal). Rectal temperature, milk output (by weight), and feed intake were measured twice daily. Mastitis, a defining feature of both natural and experimental IAV(H5N1) infection^19^, was monitored daily.

Following inoculation, all previously infected cows rechallenged with the homologous B3.13 virus remained apparently healthy, maintaining normal body temperature, milk production, and feed intake throughout the study (**Figure 2A** and **2B**). Additionally, all homologous rechallenge cows, regardless of dose, remained free of mastitis throughout the study period with no changes in milk color or consistency (**Supplemental Table 2**).

**Figure 2.**
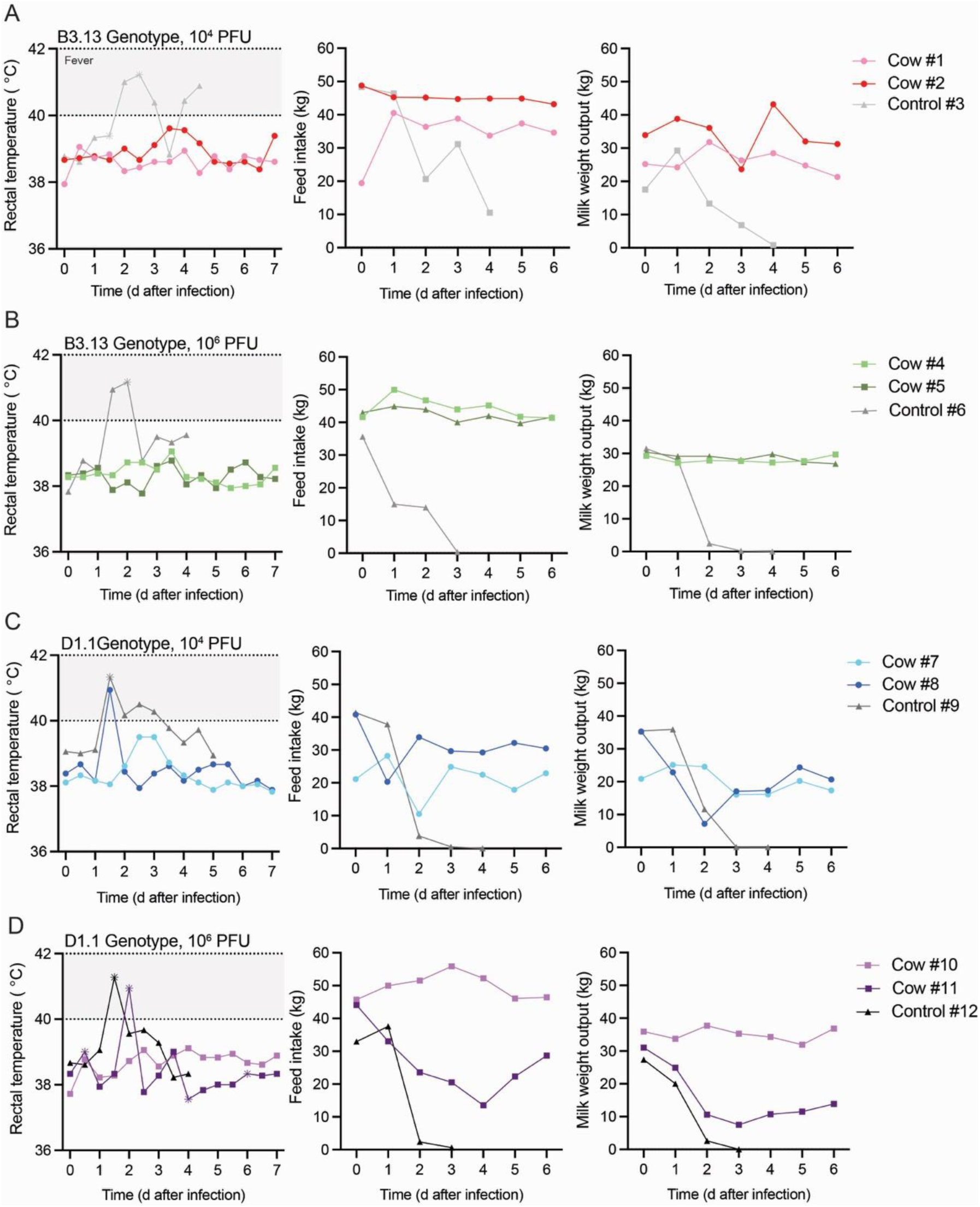
Naturally acquired IAV(H5N1) immunity protects against severe morbidity and mortality. Rectal temperature was measured twice daily and total feed intake and milk output (by weight) were reported daily for cows inoculated with (**A**) B3.13 genotype at 10⁴ PFU, (**B**) B3.13 genotype at 10⁶ PFU, (**C**) D1.1 genotype at 10⁴ PFU, and (**D**) D1.1 genotype at 10⁶ PFU. Each infection group included two previously exposed cows and one naïve control cow for comparison. None of the previously exposed cows developed severe clinical disease, whereas all control cows exhibited severe clinical signs that met endpoint criteria necessitating premature euthanasia. Control cow #3, #6, and #12 were euthanized on 4 dpi, while control cow #9 was euthanized at 5 dpi. Asterisks (*) indicate administration of flunixin meglumine (a non-steroidal anti-inflammatory drug), which corresponds with observed drops in temperature.

In contrast to the homologous virus challenge, two of the previously infected cows (cow #8 and #11) rechallenged with a heterologous D1.1 genotype virus developed transient fevers 1-2 days post-inoculation (dpi), accompanied by corresponding reductions in feed intake and milk yield (**Figure 2C** and **2D**). Both cows showed signs of recovery by the end of the study period.

More specifically, cow #8 (D1.1 genotype, 10^4^ PFU) exhibited a self-limiting fever by 1 dpi that coincided with a 20 kg reduction in daily feed intake, both of which resolved within 24 hours without treatment. Milk output declined by 28 kg at 2 dpi and began to recover by 3 dpi, though it did not fully return to prechallenge milk production (**Figure 2C**). Cow #11 (D1.1 genotype, 10^6^ PFU) exhibited moderate clinical disease, including a fever (40.9 °C) by 2 dpi. Daily feed intake and milk output decreased by 30 kg and 24 kg, respectively, with only gradual recovery by 4-5 dpi (**Figure 2D**). It should be noted that cow #11 arrived at the BSL-3Ag facility with an interdigital erosive ulcer on her left rear foot. The attending veterinarian prescribed a nonsteroidal anti-inflammatory drug and a broad-spectrum antibiotic for this lesion, which may have affected the observed clinical signs. Additionally, cow #7 (D1.1 genotype, 10^4^ PFU/cow) exhibited a slight increase in rectal temperature and brief reductions in feed intake and milk output at 2 dpi, but these parameters stabilized to levels observed prechallenge by 3 dpi (**Figure 2C**).

The clinical observations in rechallenge animals contrast sharply with those of naïve control cows (**Figure 2**), which suffered severe disease. All naïve control cows developed fever by 1-2 dpi and a nearly complete loss in feed intake and milk yield across both genotypes and doses. These severe clinical signs necessitated euthanasia at 4-5 dpi upon meeting endpoint criteria. Control cow #3 (B3.13 genotype, 10^4^ PFU) presented with a lesion on her RF foot while in the BSL-3Ag facility, which was treated with a nonsteroidal anti-inflammatory drug and a broad-spectrum antibiotic. Additionally, control cows #9 (D1.1 genotype, 10^4^ PFU) and #12 (D1.1 genotype, 10^6^ PFU) entered the BSL-3Ag facility with subclinical mastitis, which was detected during the first milk evaluation (**Supplemental Table 2**). It is uncertain whether these co-morbidities contributed to the severity of the disease after IAV(H5N1) inoculation. However, control cows #9 and #12 were clinically normal upon entry, and the subclinical mastitis would likely have gone unnoticed and untreated on-farm. Because subclinical mastitis is common in dairy herds^20,21^, these cows served as appropriate controls resembling typical farm conditions.

Altogether, these findings demonstrate that prior exposure to IAV(H5N1) provided strong protection against re-challenge with the homologous B3.13 virus, while only partial, but clinically meaningful, protection was observed following challenge with a heterologous D1.1 virus. By comparison, naïve control cows quickly succumbed to severe clinical disease, underscoring how prior immunity dramatically altered disease outcomes.

### IAV(H5N1) viral RNA was detected in milk from previously infected cows, with limited to no recovery of infectious virus

We next evaluated whether prior exposure to IAV(H5N1) limits viral shedding in milk. Following uniform virus inoculation into each mammary gland quarter, milk was collected daily from each quarter and assessed for viral RNA by rRT-PCR. Cows rechallenged with either the homologous B3.13 virus (**Figure 3A** and **3B**) or the heterologous D1.1 virus (**Figure 3D** and **3E**) exhibited markedly reduced viral RNA levels compared with the naïve controls.

**Figure 3.**
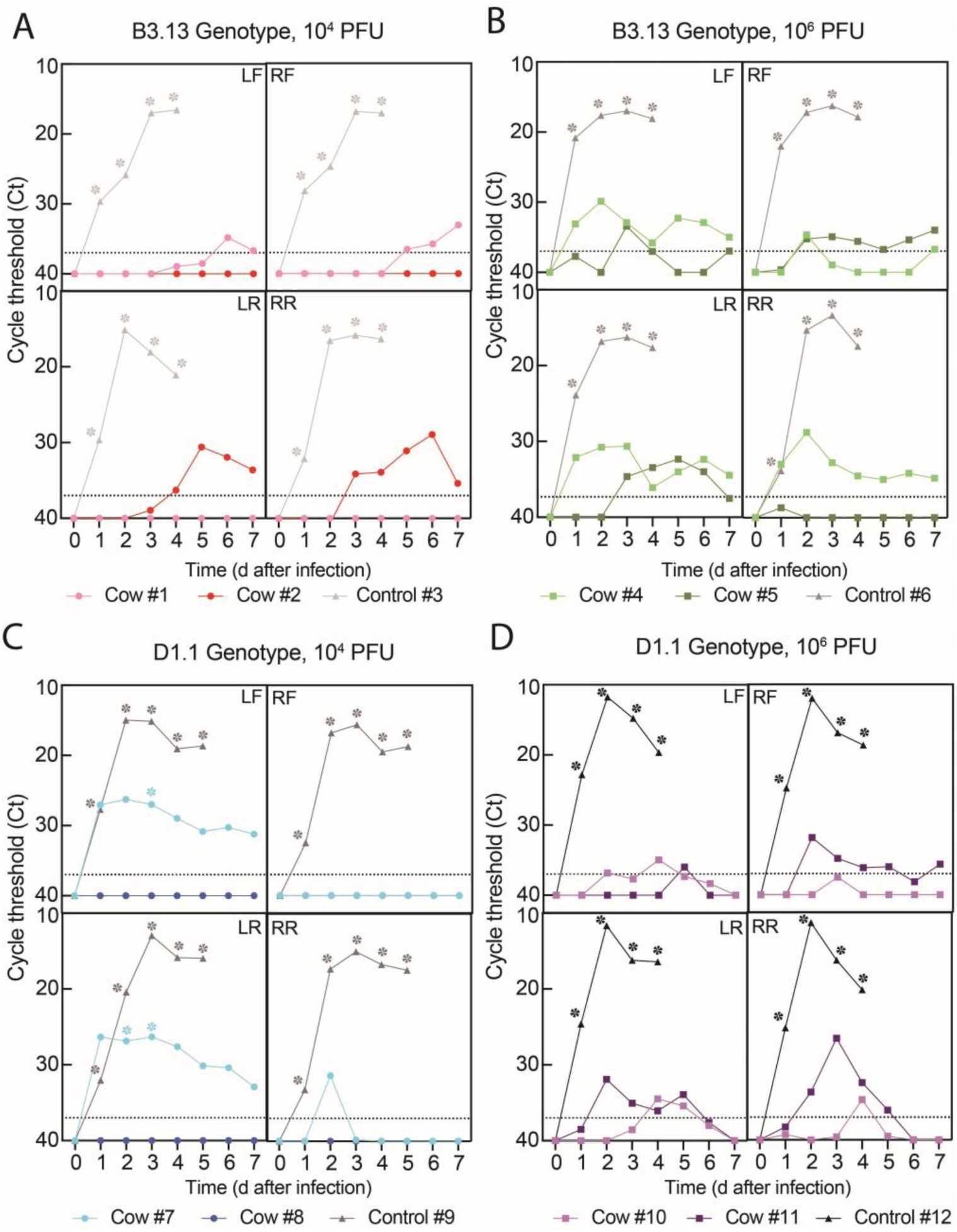
IAV(H5N1) viral RNA detected in milk from previously infected cows, with limited recovery of infectious virus. Cycle threshold (Ct) values were measured by rRT-PCR from milk collected daily for 7 days post-inoculation. Infectious virus production was quantified using TCID_50_ assays for milk samples with a Ct <32. Data are shown separately for each udder quarter: left front (LF), right front (RF), left rear (LR), and right rear (RR). Viral RNA in milk from cows inoculated with (**A**) B3.13 genotype at 10^4^ PFU, (**B**) B3.13 genotype at 10^6^ PFU, (**C**) D1.1 genotype at 10^4^ PFU, and (**D**) D1.1 genotype at 10^6^ PFU. The dotted line indicates the assay limit of detection. Asterisks (*) indicate samples in which infectious virus was recovered in milk.

In the homologous B3.13, 10^4^ PFU infection group (**Figure 3A**), viral RNA was detected in milk from only two quarters for both cow #1 (LF, RF) and cow #2 (LR, RR). Instead, in the homologous B3.13, 10^6^ PFU infection group (**Figure 3B**), viral RNA was predominantly detected in milk from three quarters for cow #4 (LF, LR, RR) and cow #5 (LF, LR, RF). Despite the detection of viral RNA, no infectious virus was recovered from cows rechallenged with the homologous B3.13 virus. These findings demonstrate clear quarter-dependent viral shedding in cows with natural immunity to IAV(H5N1), driven largely by quarter-specific differences in local immunity.

In the heterologous D1.1, 10^4^ PFU infection group (**Figure 3D**), cow #7 exhibited the lowest Ct values (Ct 26) of the previously infected cows, confined to two udder quarters (LF and LR). Infectious virus was isolated from these quarters at 2 and 3 dpi (denoted with “*” in **Figure 3D**). The infectious titers (1.0×10^3^-1.5×10^4^ TCID_50_/mL) were markedly lower than those observed from naïve control cows. Importantly, these same quarters exhibited the lowest milk virus neutralization titers before rechallenge **Figure 1B**)—both at the assay’s limit of detection—and subsequently developed clinical mastitis (**Supplemental Table 2**). In contrast, the other quarters from cow #7 (RF and RR), from which no viral RNA and no infectious virus was recovered, had higher virus neutralization titers and did not develop mastitis. Of note, no viral RNA was detected in the milk from any quarter of cow #8 (D1.1, 10⁴ PFU) (**Figure 3D**), which also exhibited the highest milk neutralization titers (all ≥ 32) (**Figure 1B**). The complete absence of detectable viral RNA or infectious virus across all four quarters strongly suggests that a neutralization titer of 32 may represent a biologically meaningful correlate of protection.

Although this neutralizing antibody titer appeared to confer sterilizing immunity, cow #8 nonetheless developed mastitis in three quarters (**Supplemental Table 2**), indicating that inflammatory processes can still arise in the absence of viral replication. Similarly, cow #10 in the heterologous D1.1, 10^6^ PFU infection group exhibited only sporadic detections of viral RNA in all four quarters, just above the limit of detection, while cow #11 of the same group had detectable viral RNA in three quarters (LR, RR, RF). Collectively, these findings indicate that prior exposure to IAV(H5N1) substantially limits viral replication but does not induce sterilizing immunity.

Despite the presence of viral RNA in milk from some mammary glands, all homologous rechallenge cows were fully protected from mastitis. Conversely, infectious virus was detected in only one heterologous rechallenge cow; however, all cows in this group developed mastitis by 3 dpi, which varied by quarter, with some affected quarters exhibiting gross abnormalities in milk color and consistency (**Supplemental Table 2**). Mastitis was consistently present in quarters that exhibited higher viral RNA (**Figure 3D** and **3E**) and lower milk neutralization titers (**Figure 1B**). This underscores the complex interplay between viral replication, local immunity, and inflammation in the development of mastitis during IAV(H5N1) infection. By comparison, all control cows infected with either the B3.13 or D1.1 genotype viruses exhibited high viral loads across all four quarters by 1 dpi, reaching peak infectious titers of 1×10^8^ TCID_50_/mL (**Supplemental Figure 2A** and **2B**). These high titers were accompanied by changes in milk color, consistency, and the development of mastitis in all four quarters by 3 dpi (**Supplemental Table 2**).

Viral shedding was also monitored in nasal swabs collected daily from all animals in this study. No viral RNA was detected in any nasal swabs collected from previously infected cows, while low-level RNA (Ct >30), peaking at 3-4 dpi, was detected in the nasal swabs from the naive control cows (**Supplemental Figure 3**). Viral RNA was higher, as evident by the lower Ct values, and detected across more consecutive days in naïve control cows inoculated with the D1.1 genotype, than those inoculated with the B3.13 genotype virus. Despite this, no infectious virus was recovered from any nasal swabs collected from naïve control cows.

### Prior immunity to IAV(H5N1) restricts viral replication in the mammary gland but may induce immune-mediated pathology upon heterologous re-exposure

At 7 dpi, previously infected cows were euthanized and necropsied to assess udder pathology, while naïve control cows developed severe disease and required euthanasia at 4–5 dpi, after which necropsy was also performed. At the time of euthanasia, across all previously infected cows—regardless of genotype (B3.13 or D1.1) or infectious dose (10^4^ or 10^6^ PFU)—viral RNA in mammary tissue was detected at low levels (with most Ct values >30), and no infectious virus was isolated (**Figure 4A, 4B, 4D, 4E**). In contrast, naïve control cows exhibited uniformly high viral loads and infectious titers in the mammary gland, peaking at 10^6^ TCID₅₀/mL (**Figure 4C** and **4F**), whereas titers in the supramammary lymph node were often several logs lower. Despite the high viral loads, mammary tissues from naïve control cows appeared grossly normal, except for thickened, discolored milk within the mammary gland, findings consistent with severe mastitis.

**Figure 4.**
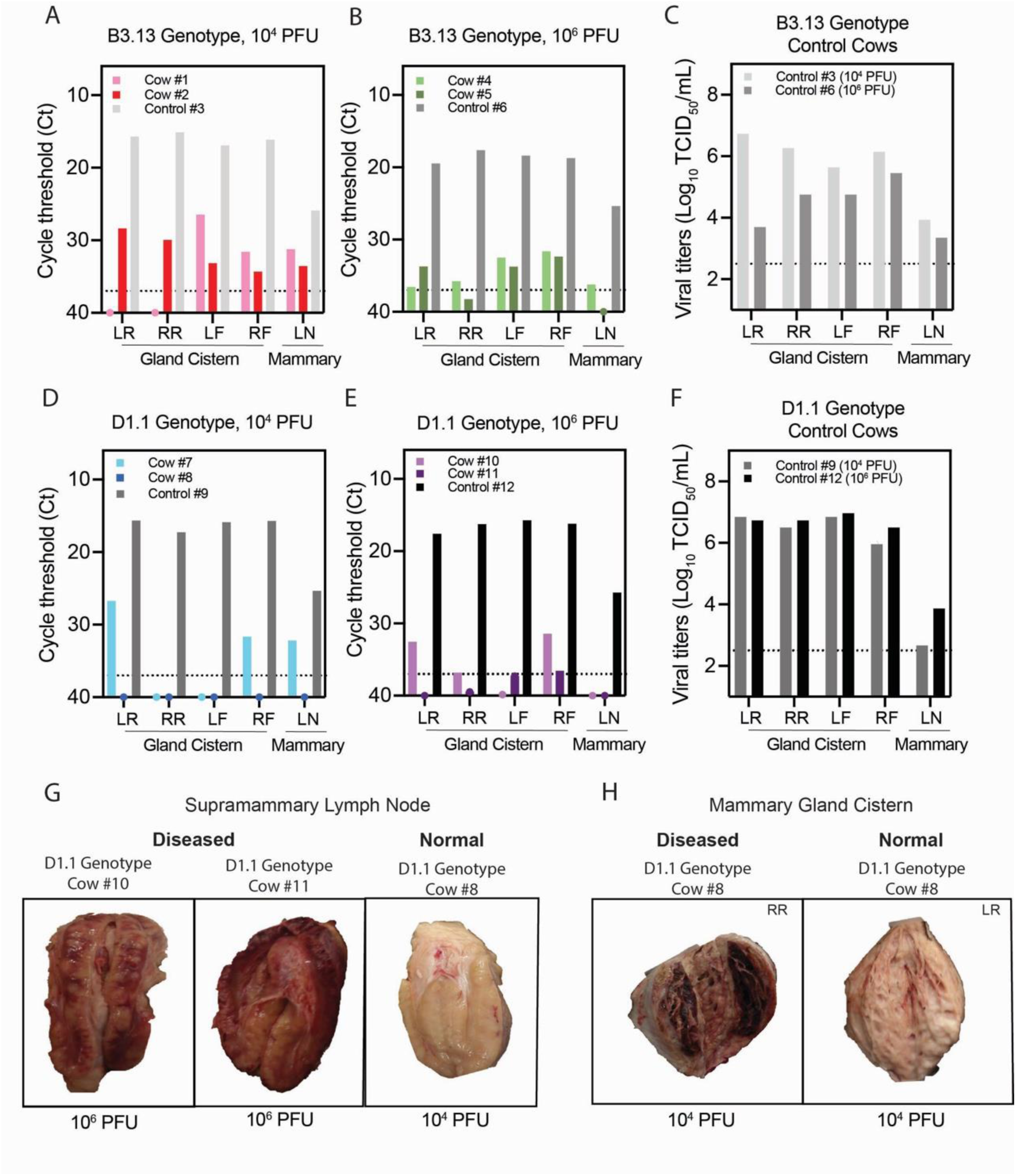
Prior immunity to IAV(H5N1) restricts viral replication in the mammary gland but may induce immune-mediated pathology upon heterologous re-exposure. Cycle threshold (Ct) values were measured by rRT-PCR from mammary tissue at necropsy conducted day 7 post-inoculation. Infectious virus production was quantified using TCID_50_ assays in tissues with a Ct <32. Data are shown separately for the gland cistern of each udder quarter—the left front (LF), right front (RF), left rear (LR), and right rear (RR)—and the supramammary lymph node (LN). (**A**) Viral RNA in tissue from cows inoculated with B3.13 genotype at 10^4^ PFU. (**B**) Viral RNA in tissue from cows inoculated with B3.13 genotype at 10^6^ PFU. (**C**) Infectious virus in tissue from control cows inoculated with B3.13 genotype. (**D**) Viral RNA in tissue from cows inoculated with D1.1 genotype at 10^4^ PFU. (**E**) Viral RNA in tissue from cows inoculated with D1.1 genotype at 10^6^ PFU. (**F**) Infectious virus in tissue for control cows inoculated with D1.1 genotype. The dotted line indicates the assay limit of detection. (**G**) Gross pathology of the supramammary lymph node, contrasting diseased tissue in the 10^6^ PFU heterologous rechallenge cows to normal tissue in the 10^4^ PFU heterologous rechallenge cows. (**H**) Gross pathology of the gland cistern in the mammary gland of cow #8, contrasting diseased tissue in the RR versus normal tissue in the LR.

Gross examination of previously infected cows revealed supramammary lymph node pathology in the heterologous D1.1, 10^6^ PFU infection group (cow #10 and cow #11) (**Figure 4G**), but not in the B3.13 rechallenge group. Viral RNA was not detected in these lymph nodes (**Figure 4E**), suggesting immune-mediated pathology rather than damage induced by viral replication. Among previously infected cows, mammary glands appeared grossly normal in nearly all quarters. The only exception was cow #8 (D1.1, 10^4^ PFU), which exhibited localized necrosis in the gland cistern of the RR quarter (**Figure 4H**). As described earlier, this cow had the highest milk neutralization titers, including a titer of 32 in the RR quarter. However, only the RR quarter was affected despite similar or higher titers (up to 64) in the other quarters (**Figure 1B**). It remains unclear whether the lesion represents sustained tissue damage from viral replication during the initial on-farm infection or immune-mediated pathology following challenge with the D1.1 genotype virus. No viral RNA was detected in mammary tissue or the supramammary lymph node of cow #8 (**Figure 4D**), which is consistent with the lack of viral shedding in milk (**Figure 3D**). For cow #7 (D1.1, 10^4^ PFU), infectious virus had been detected in milk from the LF and LR quarters (**Figure 3D**), but no infectious virus was recovered from mammary gland tissue at necropsy. The LR mammary gland contained viral RNA at levels similar to those observed in milk (Ct 26), while no viral RNA was detected in the LF mammary gland (**Figure 4D**). Overall, these findings demonstrate that heterologous viral re-exposure can drive immune-mediated pathology in cows with prior immunity to IAV(H5N1).

## DISCUSSION

Our findings demonstrate the durability of naturally acquired IAV(H5N1) immunity in dairy cows, showing strong homologous protection and clinically meaningful heterologous protection one year after the initial infection. Cows previously infected with a B3.13 genotype virus retained measurable antibodies to the infecting virus in both serum and milk more than a year later. Upon intramammary rechallenge with homologous virus, previously infected cows displayed robust protection from clinical disease, showing no fever, loss of appetite, reduction in milk output, or evidence of mastitis and no infectious virus was recovered from milk. In contrast, heterologous rechallenge induced moderate clinical signs in some cows and infectious virus was detected at low titers in the milk of one cow, indicating partial protection. By comparison, naïve control cows developed severe disease requiring euthanasia by 4–5 dpi, with high viral loads shed in milk from all four quarters. These findings are promising for future vaccination strategies yet also underscore that viral evolution may permit breakthrough viral shedding and potential silent transmission in dairy herds.

The route of transmission of IAV(H5N1) in dairy cows remains a key unresolved question in the outbreak. Both experimental and on-farm investigations demonstrate a profound tropism of the virus for the mammary gland, yielding high titers of infectious virus in milk^4,15–17,22,23^. Accordingly, the prevailing hypothesis is that transmission occurs via contaminated milking equipment^24,25^, leading to intramammary infection. While plausible, this hypothesis faces two major challenges. First, intramammary transmission (including the use of shared milking equipment) has not yet been replicated under experimental conditions^23^. Second, it does not directly address how the virus initially spread from wild birds to cattle, nor for the seroconversion observed in dry, non-lactating cows^17,26,18^, suggesting that exposure is likely multifaceted^27^. Nonetheless, more than one year later, these previously infected cows exhibited detectable serum antibodies and quarter-specific antibody responses in milk. This variation in milk antibody responses indicates a heterogeneous exposure and immune response across the udder, with potential implications for quarter-specific viral replication and shedding following re-infection. These findings underscore the functional independence of each quarter, capable of mounting a distinct local immune response. Interestingly, a previous study demonstrated that cows inoculated in only the hindquarters produced neutralizing antibodies in milk from all four quarters, protecting against clinical disease and viral replication^22^. However, antibody titers remained highest in the inoculated quarters, indicating that while systemic responses may contribute to protection—likely through the transudation of serum antibodies into milk—local immunity is also induced. Together, these observations raise important questions about the levels and localization of neutralizing antibodies required to prevent both clinical disease and viral shedding within the mammary gland. They also underscore the potential advantage of systemic vaccination, which can generate more uniform immunity, over the heterogenous immune response across the udder derived from natural exposure. Moreover, the mechanisms by which different exposure routes stimulate serum versus milk neutralizing antibodies remain poorly defined. Addressing these knowledge gaps will require further investigation into alternative routes of infection and the distinct immune responses they elicit.

The bovine mammary gland is anatomically compartmentalized into four distinct quarters, each capable of becoming independently infected with IAV(H5N1) without evidence of inter-quarter spread^23^. This functional independence was further supported in our study by the heterogeneous distribution of neutralizing antibodies across quarters, raising questions about exposure in these animals and whether this uneven quarter-level immunity may permit breakthrough viral shedding in some quarters. As a result, herds with prior immunity may act as silent spreaders of IAV(H5N1), remaining protected from clinical disease while still permitting low-level viral shedding. Here—across both viral genotypes and infectious doses—viral RNA was detected in milk at low levels in previously infected animals compared with naïve control cows. No infectious virus was recovered from homologous rechallenge animals, whereas low infectious titers were detected at 2–3 dpi in one heterologous rechallenge cow. All previously infected cows rechallenged with either homologous or heterologous virus exhibited quarter-dependent shedding of viral RNA. This heterogeneous pattern—also noted in on-farm investigations^27^—is likely driven by the varying immune protection among quarters, consistent with the antibody profiles observed here. Further, homologous rechallenge cows did not develop mastitis in any quarter, whereas heterologous rechallenge cows showed quarter-specific mastitis that aligned with their patterns of viral shedding, supporting incomplete protection following heterologous re-exposure. In contrast, naïve control cows inoculated uniformly in all four quarters developed mastitis and shed high titers of infectious virus from each inoculated quarter. While necessary for comparison, the naïve control cows were obtained from a different herd than the naturally immune cows. As a result, herd-level differences in microbiome composition, genetics, management, and nutrition, may have influenced immune profiles and viral kinetics observed here. Nevertheless, these findings suggest that re-exposure to a homologous virus would substantially limit transmission in dairy herds, as viral replication remains highly restricted compared to control cows, and no infectious virus is shed. In contrast, introduction of a heterologous virus could permit breakthrough shedding, enabling the virus to persist and silently spread within the herd.

Severe morbidity and mortality associated with IAV(H5N1) infection have been documented on-farm, resulting in substantial economic losses^17^. These outcomes have also been reproduced under experimental conditions, underscoring the considerable burden this virus poses to the U.S. dairy industry. Here, experimental rechallenge of cows previously infected with IAV(H5N1) B3.13 genotype demonstrated robust protection from clinical disease upon homologous rechallenge, indicating a durable immune response persisting more than one year later. In contrast, rechallenge with a heterologous virus resulted in only partial protection, with some cows experiencing transient fever, reductions in feed intake, and milk yield. While these cows did not develop the severe clinical disease observed in naïve controls, their milk production losses were still considerable and would likely have economic consequences in natural settings. Given the short study period, additional studies are needed to determine whether milk yield recovers sufficiently and promptly for cows to remain in the herd, as well as to evaluate the economic implications. Together, these findings indicate that prior exposure confers robust protection against homologous virus and meaningful cross-protection against heterologous virus. The longevity of this protective immunity remains to be determined and has important implications for the trajectory of the IAV(H5N1) outbreak in the U.S. dairy herd. Further, the introduction of immunologically naïve animals through herd replacement will alter the immune landscape, supporting the rationale for future vaccination strategies.

Early in the outbreak, reported cases of IAV(H5N1) in dairy herds were limited. Because IAV(H5N1) infection was unprecedented in dairy cows, many cases likely went undetected, as suggested by findings from retail milk surveillance^28^. Following the initiation of federal surveillance programs^29^, case detections increased rapidly and it was noted that IAV(H5N1) was distributed across herds nationwide. More recently, new detections have declined, with only two new reported cases in the past 30 days^13^. This is likely influenced by substantial herd immunity resulting from widespread exposure of dairy cattle over the past few years. Although this has positive implications for reducing the burden of disease on the dairy industry, ongoing monitoring of viral evolution in wild birds remains essential, as continued adaptation could lead to new introductions into cattle, including genotypes with altered antigenicity that may evade existing immunity. Similarly, commercial and backyard poultry flocks remain heavily impacted by IAV(H5N1)^30^, which underscores an additional risk of spillover into dairy herds.

Because the IAV(H5N1) D1.1 genotype virus spilled over into dairy cows in early 2025 following the initial B3.13 outbreak, its pathogenesis in dairy cows remains poorly defined under experimental conditions. Naïve control cows inoculated intramammary with either the B3.13 or D1.1 genotype at both infectious doses (10^4^ or 10^6^ PFU/cow) developed comparable clinical disease, including fever by 1–2 dpi accompanied by marked reductions in feed intake and milk production, leading to humane euthanasia by 4–5 dpi. Replication kinetics were also similar between genotypes in control cows, with all four quarters exhibiting high viral loads and infectious titers exceeding 1×10^8^ TCID_50_/mL. Naïve control cows inoculated with 10^4^ PFU showed delayed replication kinetics; however, peak titers still surpassed 1×10^8^ TCID_50_/mL in milk. These findings indicate that the pathogenesis and susceptibility of dairy cows to IAV(H5N1) B3.13 genotype and D1.1 are broadly comparable. At necropsy, mammary glands from most previously infected cows rechallenged with either genotype or infectious dose appeared grossly normal. The exception was cow #8 (D1.1 genotype, 10^4^ PFU), which exhibited localized necrosis within the gland cistern of the RR quarter only. Upon arrival to the facility and at the initial milking, the RR quarter of this cow was firm on palpation after milking. Previous studies have reported that IAV(H5N1) infection may have long-term impacts on dairy cows as a consequence of the virus-induced mastitis and damage to milk-producing epithelial cells^17^, though we cannot determine whether this damage resulted from the initial infection. No viral RNA was detected in the milk from any quarter of this cow after heterologous challenge, indicating sterilizing immunity. The restricted replication observed in this cow and other previously infected animals reflects immune-mediated control by neutralizing antibodies, but a reduced number of functional epithelial cells remaining after virus-induced damage during the initial infection may also play a role. Beyond the mammary gland, cows rechallenged with the heterologous D1.1 genotype at the high infectious dose (10^6^ PFU) showed marked hemorrhage within the supramammary lymph node, likely reflecting an exaggerated local immune response. Consistent with this immune-mediated pathology, only limited viral RNA was detected in mammary gland tissue and the supramammary lymph node of previously infected cows, and infectious virus was not recovered, in stark contrast to naïve control cows. The B3.13 genotype and D1.1 genotype share highly similar HA sequences but differ more substantially by their internal gene segments. The B3.13 genotype emerged after a European-origin 2.3.4.4b entered North America in 2021, and acquired PB2, PB1, NP and NS segments via reassortment with local LPAI viruses^18,26,31,32^. Instead, the D1.1 genotype had acquired the PB2, PA, NP and NA segments from local LPAI viruses^32,33^. Internal gene segments of highly pathogenic avian influenza viruses are known to influence viral replication dynamics and host immune responses^34–36^. Accordingly, the divergent pathology observed following homologous versus heterologous rechallenge likely reflect a combination of viral genetic differences—including internal gene segments and antigenic variation in HA and NA—as well as immune factors beyond neutralizing antibodies, such as cellular immunity, which warrant further investigation.

As IAV(H5N1) spreads among dairy cows, there is growing concern that these viruses could acquire mammalian adaptations that enhance their ability to infect humans. Signatures of mammalian adaptation have already been detected in IAV(H5N1) viruses from dairy cows, including markers associated with increased replication and transmission efficiency^32,37^. Recent data show that viruses of the B3.13 and D1.1 genotypes replicate more efficiently in bovine cells than other avian- or mammalian-origin IAVs^37^, raising questions about the broader susceptibility of dairy cattle to diverse influenza viruses. Although this susceptibility remains poorly defined, the extensive genomic diversity of IAV(H5N1) viruses circulating in wild birds—combined with close interactions among humans and other livestock species such as swine that maintain endemic IAVs—creates ongoing opportunities for spillover and reassortment in dairy cows. Our findings of breakthrough shedding of infectious virus in a heterologous rechallenge cow, despite only mild clinical signs and limited tissue pathology, underscore the importance of studies that investigate the susceptibility of dairy cows to diverse IAVs, given the potential for silent spread. However, prior infection with the IAV(H5N1) B3.13 genotype provided substantial protection against severe disease and restricted viral replication following both homologous and heterologous challenge. These data suggest that vaccination strategies inducing robust mucosal immunity could serve as a promising intervention to reduce additional spillover opportunities, virus spread, and disease burden within U.S. dairy herds. Further, our results highlight the importance of including multiple circulating genotypes and/or drift variants in future vaccine regimens given the potential for milk production losses, breakthrough infections, and tissue damage observed in the heterologous rechallenge group. Additional work is needed to determine the precise correlates of protection in dairy cattle exposed to IAV (H5N1). It is unclear whether immune-mediated protection is driven primarily by humoral responses (e.g., could passively administered convalescent sera serve as a potential therapeutic) or whether tissue-resident T cells within the mammary gland also plays an essential role in viral control.

## MATERIALS AND METHODS

### Research ethics and safety

All animal experiments conducted at The Ohio State University were approved by the Institutional Animal Care and Use Committee of the Ohio State University under protocol #2025A00000035. IAV(H5N1) research was approved by The Ohio State University Institutional biosafety committee under approval #2025R00000037 and all virus experiments were performed in The Ohio State University Ralph Regula Plant and Animal Agricultural Research (PAAR) Biosafety Level 3 (BSL3) large animal facility in Wooster, OH, USA.

### Animals, study design, and sample collection

In April 2024, an outbreak of IAV(H5N1) occurred in an Ohio dairy herd, during which individual-level diagnostic testing was performed on selected clinically ill animals. One year later, in April 2025, we identified eight cows remaining in the herd that had been clinically affected during the outbreak and confirmed to be infected with the B3.13 genotype through rRT-PCR testing of composite milk samples representing each mammary gland quarter collected on April 5, 2024. For comparison, four naïve control cows were sourced from a separate herd with no documented exposure to IAV(H5N1). Prior to rechallenge, all previously infected cows had detectable antibodies to IAV(H5N1), while control cows were negative for antibodies in serum and milk (**Figure 1A and B**). Following antibody characterization, cows were assigned to four groups receiving either B3.13 or D1.1 at 10^4^ or 10^6^ PFU per cow. Using two inoculation doses (10^4^ and 10^6^ PFU), we could evaluate whether prior immunity modulated disease severity, viral replication kinetics, or antibody responses in a dose-dependent manner, with the higher dose representing a previously characterized lethal challenge dose^16^. Before inoculation, teats were aseptically cleaned with povidone-iodine and 70% ethanol. All animals were inoculated intramammary using a sterile teat cannula (Jorgensen Laboratories, J-12 Teat Infusion Cannula). Viral stocks (e.g., 10^4^ or 10^6^ PFU/cow) were first resuspended in 10 mL of infection media (Eagle’s Minimum Essential Medium (EMEM)) and then distributed evenly across all four quarters (2.5 mL per teat canal). Following inoculation, teats were gently massaged to facilitate upward distribution of the inoculum. Cows were milked twice daily using a bucket milker. Before the morning milking, milk samples were collected separately from each udder quarter. To prevent cross-contamination, gloves were changed between quarters. Mastitis was assessed once daily using the California Mastitis Test (CMT). Clinical monitoring included twice-daily measurement of rectal temperature, quantification of milk output by weight, and assessment of feed consumption and refusal. Animals were euthanized and necropsied at 7 days post-inoculation or earlier if predefined humane endpoints were reached.

### Virus isolation and propagation

The homologous A/bovine/Ohio/B24OSU-342/2024 (H5N1) virus (GenBank IDs: PP836412.1–PP836419.1), which originated from the same Ohio dairy farm where we sourced our eight previously infected cows, and the heterologous A/bovine/Nevada/WD-210/2025 (H5N1) virus (GenBank IDs: PV520600.1–PV520607.1) were isolated from milk samples collected during dairy cattle outbreaks in Ohio and Nevada, respectively. Both viruses were initially amplified by two serial passages in 10-day-old embryonated chicken eggs, followed by one (D1.1) or two (B3.13) low multiplicity of infection (MOI) passages in Madin-Darby canine kidney (MDCK) cells. The resulting virus stocks were used for all subsequent experiments.

### Plaque assay titration

Plaque assays were performed as described previously^23^. Briefly, MDCK cells were exposed to 10-fold serial dilutions of virus in serum-free EMEM (0.8 mL per well). Cells were incubated with virus for 1 hour at 37 °C with gentle rocking every 15 minutes. After incubation, the inoculum was removed and 3 mL of overlay medium (EMEM supplemented with L-glutamine and PenStrep, 2.8% Avicel, and 1 µg/mL TPCK-treated trypsin) was added. Plates were incubated for 48 hours at 37 °C with 5% CO₂, after which the cells were washed and fixed with 20% methanol containing 0.2% crystal violet. Plates were rinsed with deionized water, plaques were enumerated, and viral titers were calculated as plaque-forming units per milliliter (PFU/mL).

### Infectious virus titration by TCID_50_ assay

Biomaterials (e.g., milk, nasal swabs) and mammary tissue homogenates with Ct values <32 were subjected to infectious virus titration using a TCID_50_ assay in MDCK cells. Samples were initially diluted 1:100 in infection medium (EMEM supplemented with 0.3% bovine serum albumin [BSA] and 1 µg/mL TPCK-treated trypsin), then serially 5-fold diluted and plated (100 µL per well) in quadruplicate onto confluent MDCK cells (5×10^4^ cells/well, plated the day prior in 96-well plates). Non-inoculated wells were included on each plate and served as negative controls. Plates were incubated for 72 hours at 37°C in 5% CO₂. After incubation, the media was removed and the cells were washed 1x with PBS, followed by a 20-minute staining with crystal violet solution (20% methanol containing 0.2% crystal violet in PBS). After thorough PBS washes, each well was scored as negative or positive for cytopathic effect (CPE), and TCID_50_ titers were calculated using the Reed–Muench method.

### RT-qPCR for IAV RNA load detection

RNA was extracted from biomaterials and tissue homogenates using the MagMAX™ Viral/Pathogen Nucleic Acid Isolation Kit (Applied Biosystems), according to the manufacturers’ protocols. Extracted RNA was tested for the presence of viral RNA using rRT-PCR with the VetMAX™-Gold SIV Detection Kit (Applied Biosystems). Reactions were performed on a QuantStudio™ 5 Real-Time PCR System (Applied Biosystems).

### Virus neutralization assay

Antibodies in serum and milk collected separately for each udder quarter— left front (LF), right front (RF), left rear (LR), and right rear (RR)— in cows prior to experimental re-infection were quantified by a virus neutralization assay using MDCK cells. Milk samples were treated with Rennet from *Mucor miehei* (MilliporeSigma, R5876) at a concentration of 5 mg/mL until curdled, then centrifuged at 10,000 rpm for 5 min to remove fat and casein. The clarified liquid layer was collected. Serum samples were heat-inactivated at 56°C for 30 min and diluted 1:10 in infection medium (EMEM supplemented with 0.3% bovine serum albumin [BSA] and 1 µg/mL TPCK-treated trypsin). Milk samples were heat-inactivated at 60°C for 30 min and diluted 1:2 in infection medium. All samples were then serially 2-fold diluted and incubated with 100 TCID₅₀ of IAV(H5N1) B3.13 genotype for 1 h at room temperature. Virus–sample mixtures were then added in quadruplicate to MDCK cell plates and incubated for 72 hours at 37°C in 5% CO₂. After 72 h, cells were rinsed with PBS, fixed, and stained with crystal violet to assess cytopathic effect (CPE). The virus neutralization titer was defined as the highest dilution protecting at least 2 out of 4 replicate wells from CPE.

### Hemagglutination inhibition assay

Serum samples were treated with receptor-destroying enzyme (RDE) and incubated at 37°C for 18–20 h, followed by heat inactivation at 56°C for 30 min. Treated serum was serially 2-fold diluted and mixed with 4 HAU (hemagglutination units) of an attenuated rg-A/American wigeon/South Carolina/22-000345-001/2021 (H5N1) (HA modified, NA)+ PR8 [R] (6+2) virus for 1 h at room temperature. Agglutination was subsequently assessed using 1% horse red blood cells, and the HAI titer was defined as the highest serum dilution completely inhibiting hemagglutination, performed in duplicate.

### Tissue homogenization

Frozen tissue samples were thawed at room temperature and transferred to TissueLyser II (Qiagen) homogenization cassettes. Samples were homogenized at 20 Hz for 60 seconds, followed by 30 Hz for 60 seconds. The cassettes were then flipped, and the homogenization cycle was repeated to ensure complete tissue disruption. The samples were then centrifuged at 12,000 rpm for 1-minute, and the supernatant was aliquoted for subsequent assays.

## DATA AVAILABILITY

The authors declare that all data supporting findings are available within the paper and supplemental information.

## MATERIALS AVAILABILITY

This study did not generate new unique reagents.

## ACKNOWLEDGEMENTS

We would like to thank PAAR facility staff (Alden Sewell, Kaitlynn Starr, Jonathan Zsoldos, and Michael Jeffers) for their support of this study. Thank you to Justin Kieffer and Brad Youngblood for their assistance in acquiring animals for this study and with veterinary care. We also thank John Franks, Jennifer DeBeauchamp, David Walker, Trushar Jeevan, and Tom Fabrizio for the initial isolation and characterization of the influenza A(H5N1) B3.13 and D1.1 genotype viruses. Figures supporting experimental design were created using https://BioRender.com.

Funding was provided by the Centers of Excellence for Influenza Research and Response (CEIRR), National Institute of Allergy and Infectious Diseases, National Institutes of Health, Department of Health and Human Services under contract 75N93021C00016.

## AUTHOR CONTRIBUTIONS

Conceptualization-RJW, CJW, ASB

Methodology-NNT, RJW, CJW, ASB

Data curation-all

Visualization-NNT, CJW, ASB

Animal care-NNT, HJC, EAM, ML, KSK, CJW, ASB

Writing, original draft-NNT, CJW, ASB

Writing, editing/review-all

## SUPPLEMENTAL INFORMATION INDEX

Supplemental Figures 1-3

Supplemental Tables 1 and 2

