## Supplemental for "Natural H5N1 immunity in dairy cows is durable and cross-protective but non-sterilizing"

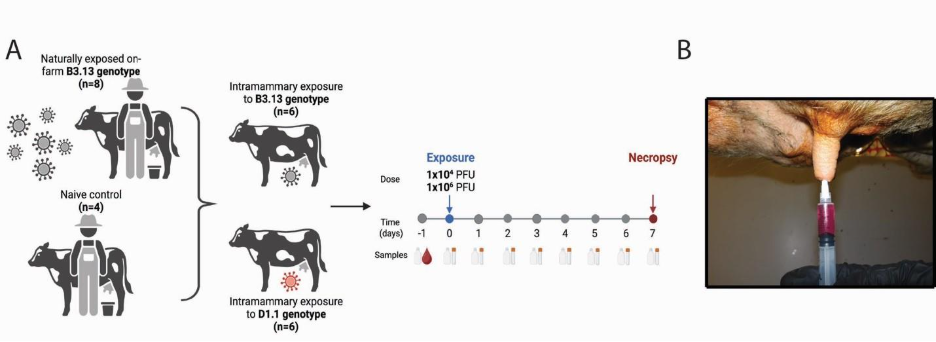

**Supplemental Figure 1. Schematic of experimental design (related to Figures 2 and 3).** (**A**) Cows naturally exposed on-farm to IAV(H5N1) B3.13 genotype (n = 8) or naïve controls obtained from a different unexposed farm (n = 4) were subsequently rechallenged with either homologous B3.13 genotype (n = 6) or heterologous D1.1 genotype (n = 6) viruses. (**B**) Cows were inoculated intramammary using a teat cannula with either 10^4^ or 10^6^ PFU/cow. A 10 mL suspension of viral inoculum was prepared for each cow and divided equally among the four teats (2.5 mL per teat).

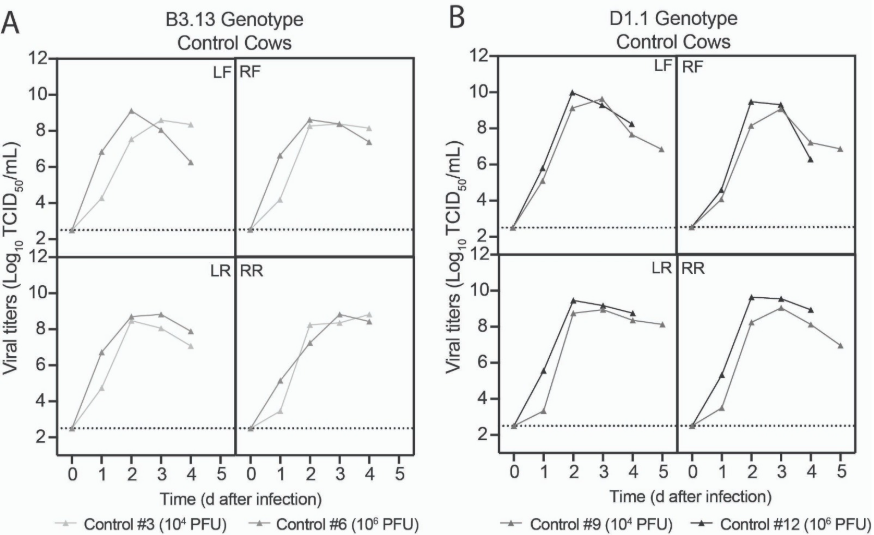

**Supplemental Figure 2. Control cows displayed robust recovery of infectious virus in milk across all four quarters (related to Figure 3).** Data are shown separately for each udder quarter: left front (LF), right front (RF), left rear (LR), and right rear (RR). Infectious virus production was quantified using TCID_50_ assays in milk from control cows inoculated with the (**A**) B3.13 genotype virus and (**B**) D1.1 genotype virus.

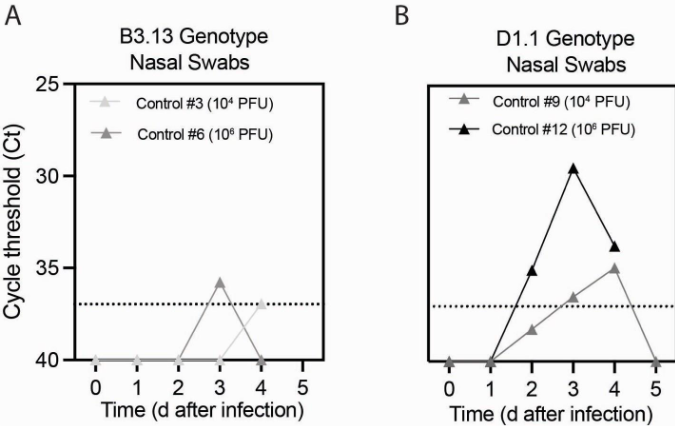

**Supplemental Figure 3. Detection of low-level viral RNA in nasal swabs of naïve control cows, with no recovery of infectious virus (related to Figure 3).** Viral RNA was quantified by rRT-PCR from nasal swabs collected daily from control cows inoculated intramammary with either (**A**) B3.13 genotype or (**B**) D1.1 genotype at the indicated infectious doses.

**Supplemental Table 1 (related to Figure 1).** rRT-PCR cycle threshold (Ct) values from composite milk samples representing each mammary gland quarter collected on April 5, 2024. These eight cows remained in the herd and exhibited clinical illness during the Ohio outbreak.

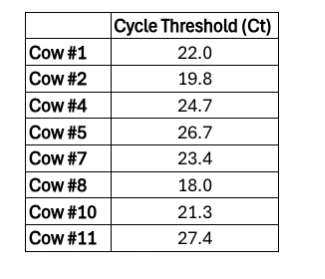

**Supplemental Table 2. Clinical mastitis results displayed daily for 7 days post-inoculation (related to Figures 1-3).** Data are shown separately for each udder quarter: left front (LF), right front (RF), left rear (LR), and right rear (RR). Mastitis status was evaluated using the California Mastitis Test, and milk changes were recorded through visual observation. Color shading indicates treatment groups: green for homologous B3.13 rechallenge, orange for heterologous D1.1 rechallenge, and gray for naïve controls. Darker shading within cells marks the onset of quarters with observed changes in milk color or consistency.

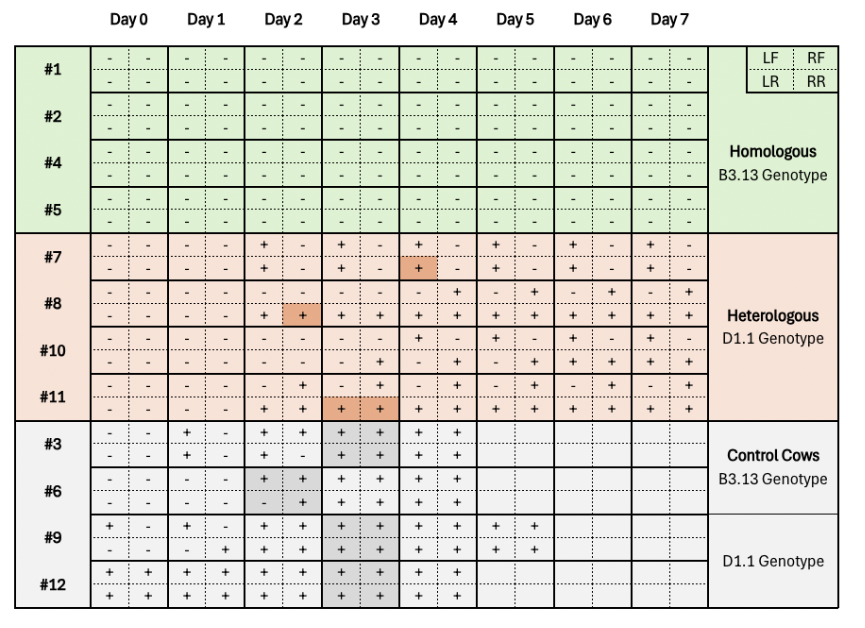
